# Identification of essential oils with activity against stationary phase *Staphylococcus aureus*

**DOI:** 10.1101/727883

**Authors:** Shuzhen Xiao, Peng Cui, Wanliang Shi, Ying Zhang

## Abstract

*Staphylococcus aureus* is the most dominant human pathogen, responsible for a variety of chronic and severe infections. There is mounting evidence that persisters are associated with treatment failure and relapse of persistent infections. While some essential oils were reported to have antimicrobial activity against growing *S. aureus*, activity of essential oils against the non-growing stationary phase *S. aureus* enriched in persisters has not been investigated. In this study, we evaluated the activity of 143 essential oils against stationary phase *S. aureus* and identified 39 essential oils (Cinnamon bark, Oregano, Thyme white, Bandit “Thieves”, Lemongrass (*Cymbopogon flexuosus*), Sandalwood oil, Health shield, Allspice, Amyris, Palmarosa, Cinnamon leaf, Clove bud, Citronella, Geranium bourbon, Marjoram, Peppermint, Lemongrass (*Cymbopogon citratus*), Cornmint, Elemi, Ho wood, Head ease, Lemon eucalyptus, *Litsea cubeba*, Myrrh, Parsley seed, Coriander oil, Dillweed, Hyssop, Neroli, Rosewood oil, Tea tree, Cajeput, Glove bud, Lavender, Sleep tight, Vetiver, *Palo santo*, Sage oil, Yarrow) at 0.5% concentration, 10 essential oils (Cinnamon bark, Oregano, Thyme white, Bandit “Thieves”, Lemongrass (*Cymbopogon flexuosus*), Sandalwood oil, Health shield, Allspice, Amyris, Palmarosa) at 0.25% concentration, and 7 essential oils (Cinnamon bark, Oregano, Thyme white, Lemongrass (*Cymbopogon flexuosus*), Allspice, Amyris, Palmarosa) at 0.125% concentration to have high activity against stationary phase *S. aureus* with no visible growth on agar plates after five-day exposure. Among the 10 essential oils which showed high activity at 0.25% concentration, 9 (Cinnamon bark, Oregano, Thyme white, Bandit “Thieves”, Lemongrass (*Cymbopogon flexuosus*), Health shield, Allspice, Palmarosa, Amyris) showed higher activity than the known persister drug tosufloxacin, while the other one (Sandalwood oil) was found to be active at a higher concentration. In Oregano essential oil drug combination studies with clinical antibiotics, Oregano plus quinolone drugs (tosufloxacin, levofloxacin, ciprofloxacin) and rifampin completely eradicated all stationary phase *S. aureus* cells, but had no apparent enhancement for linezolid, vancomycin, sulfamethoxazole, trimethoprim, azithromycin and gentamicin. Our findings may facilitate development of more effective treatment for persistent *S. aureus* infections.

## Introduction

*Staphylococcus aureus* is the leading cause of nosocomial and community-associated infections, which is responsible for a wide variety of infections that include mild superficial skin infections, osteomyelitis, implant-associated heart valve, native valve endocarditis, severe sepsis and bacteremia [1]. Although antibiotic resistance is a major problem in treatment of infections caused by *S. aureus*, drug-tolerant persisters are demonstrated to be significant contributors of chronic persistent infections and recurrent infections [2]. Clinically, infections caused by *S. aureus* such as soft tissue infections, endocarditis, osteomyelitis, prosthetic joint infections, and biofilm-related infections on indwelling device is difficult to cure with the current antibiotics, which are mainly active against the growing bacteria but have poor activity against the non-growing persisters [3]. Recently, it has been shown that a drug combination approach using drugs targeting both log phase growing bacteria and the non-growing stationary phase bacteria could more effectively eradicate a persistent urinary tract infection and a biofilm skin infection in the mouse models [4, 5]. However, the choice of persister drugs is limited at present, and treatment of persistent infections remains a challenge. Although some essential oils were found to be active against growing *S. aureus* [6-8], the activity of essential oils against non-growing stationary phase *S. aureus* has not been studied. Because activity against non-growing persisters or stationary phase bacteria correlates with in vivo activity against persistent infections in the context of drug combination in the case of uropathogenic *E. coli* and *B. burgdorferi* persistent infections [4, 9, 10], here, we evaluated a panel of 143 essential oils for their activity against stationary phase *S. aureus* as a model for activity against *S. aureus* persisters. We identified a range of highly potent essential oils with excellent activity against non-growing stationary phase *S. aureus.*

## Materials and Methods

### Bacterial strain

*S. aureus* Newman, a commonly used pan-susceptible strain isolated from a patient suffering from osteomyelitis [3] was used in this study. The strain was incubated in Tryptic Soy Broth (TSB) medium overnight to stationary phase without shaking at 37 °C, 5% CO_2_. The stationary phase *S. aureus* culture (∼10^9^ CFU/mL) was used directly without dilution for essential oil screens and drug exposure tests.

### Antibiotics and essential oils

Tosufloxacin, ciprofloxacin, levofloxacin, rifampin, linezolid, vancomycin, sulfamethoxazole, trimethoprim, azithromycin and gentamicin were purchased from Sigma-Aldrich (St. Louis, MO, USA) and dissolved in dimethyl sulfoxide (DMSO) or H_2_O to form stock solutions. All antibiotic stocks (except DMSO stocks) were filter-sterilized by 0.2 µm filter and stored at −20°C.

Commercially available essential oils were purchased from Natural Acres (MO, USA), Plant Therapy (ID, USA) and Plant Guru (NJ, USA). More information about the essential oils can be found on their websites (http://www.theplantguru.com/gc-ms-testing, http://www.planttherapy.com/essential-oils, http://naturalacresoils.com/collections/all).

DMSO-soluble essential oils were dissolved in DMSO at 5% (v/v). DMSO-insoluble essential oils were directly added to *S. aureus* cultures, then vortexed to form aqueous suspension. The 5% essential oils or aqueous suspension were further diluted into the bacterial cultures to achieve desired dilution in the following drug exposure or MIC experiments to evaluate their activity against non-growing stationary phase or growing log phase *S. aureus*.

### Screening of essential oils for their activity against stationary phase *S. aureus*

To evaluate the effect of essential oils on stationary phase bacteria, the essential oils and drugs were added to the 96-well plates containing stationary phase bacteria, leaving the first and last columns in each plate blank for control. In the primary screen, each essential oil was assayed at three concentrations: 0.5%, 0.25% and 0.125% (v/v). Tosufloxacin, ciprofloxacin, levofloxacin, rifampin, linezolid, vancomycin, sulfamethoxazole, trimethoprim, azithromycin and gentamicin were used at 50 µM as control antibiotics. The plates were incubated at 37 °C, 5% CO_2_ without shaking. After three days and five days of exposure to essential oils or drugs, the bacterial suspension was transferred to TSB plates with a 96-pin replicator to monitor the bacterial survival and regrowth after further incubation at 37 °C. All tests were run in triplicate.

### Antibiotic susceptibility test

The minimum inhibitory concentrations (MICs) were determined using microdilution method according to the CLSI guideline [11]. Essential oils were 2-fold diluted from 1% to 0.0075%. Gentamicin was 2-fold diluted from 512 µg/mL to 0.25 µg/mL as a control antibiotic. The 96-well plates were sealed and incubated at 37 °C overnight without shaking. All experiments were run in triplicate.

### Validation of active essential oils by colony forming unit (CFU) assay

The stationary phase bacteria were transferred into Eppendorf tubes. Essential oils were added at 0.25% and 0.125% concentrations. Tosufloxacin, ciprofloxacin, levofloxacin, rifampin, linezolid, vancomycin, sulfamethoxazole, trimethoprim, azithromycin and gentamicin were added to bacterial suspensions at the final concentration of 20 µM, respectively. At different time points, 100 µL bacterial suspensions were collected by centrifugation, washed and resuspended in PBS. After serial dilutions, 10 µL of each dilution was plated on TSB plate for CFU count.

### Drug combination assay on stationary phase *S. aureus*

In this study, we used Oregano as the common element to test the activity of various two-drug combinations in killing *S. aureus* Newman stationary phase cells. We evaluated tosufloxacin, levofloxacin, ciprofloxacin, rifampin, linezolid, vancomycin, sulfamethoxazole, trimethoprim, azithromycin and gentamicin at the final concentration of 5 μg/mL in combination with Oregano (0.025%). The designed drug combinations or single drug controls were added directly to stationary phase culture and CFU count was performed at different time points.

## Results

### Identification of active essential oils against stationary phase *S. aureus*

Consistent with our previous study [3], tosufloxacin was shown to have high activity against stationary phase *S. aureus*, while other clinical drugs including ciprofloxacin, levofloxacin, rifampin, linezolid, vancomycin, sulfamethoxazole, trimethoprim, azithromycin and gentamicin were not able to completely kill stationary phase *S. aureus* at 50 µM after five-day drug exposure [8]. Interestingly, after three-day exposure, 30 (Cinnamon bark, Oregano, Thyme white, Lemongrass (*Cymbopogon flexuosus*), Bandit “Thieves”, Sandalwood oil, Health shield, Allspice, Amyris, Palmarosa, Cinnamon leaf, Clove bud, Citronella, Geranium bourbon, Marjoram, Peppermint, Lemongrass (*Cymbopogon citratus*), Cornmint, Elemi, Ho wood, Head ease, Lemon eucalyptus, Litsea cubeba, Myrrh, Parsley seed, Coriander oil, Dillweed, Hyssop, Neroli, Rosewood oil), 6 (Cinnamon bark, Oregano, Thyme white, Bandit “Thieves”, Lemongrass (*Cymbopogon flexuosus*), Sandalwood oil) and 7 (Cinnamon bark, Oregano, Thyme white, Lemongrass (*Cymbopogon flexuosus*), Allspice, Amyris, Palmarosa) essential oils were found to have high activity against stationary phase *S. aureus* at 0.5%, 0.25% and 0.125% concentrations, respectively. When the drug exposure was extended to five days, additional 9 essential oils (Tea tree, Cajeput, Glove bud, Lavender, Sleep tight, Vetiver, Palo santo, Sage oil, Yarrow) and 4 essential oils (Health shield, Allspice, Amyris, Palmarosa) were found to be active at 0.5% and 0.25% concentration, respectively (Table 1). The top 10 essential oils (Cinnamon bark, Oregano, Thyme white, Lemongrass (*Cymbopogon flexuosus*), Bandit “Thieves”, Sandalwood oil, Health shield, Allspice, Amyris, Palmarosa), which showed high activity at 0.25% concentration, were used in the subsequent testing to confirm their activity in inhibiting growth of *S. aureus* in MIC test and in CFU drug exposure assay for their activity against non-growing stationary phase *S. aureus*.

**Table 1.** Effect of essential oils on stationary phase *Staphylococcus aureus*.

| EO<sup>a</sup> | Plant | Viability of bacteria after 3 or 5 days of exposure<sup>b</sup> |  |  |  |  |  |
| --- | --- | --- | --- | --- | --- | --- | --- |
|  |  | 0.5% EO<sup>a</sup> |  | 0.25% EO<sup>a</sup> |  | 0.125% EO<sup>a</sup> |  |
|  |  | 3 days | 5 days | 3 day | 5 days | 3 day | 5 days |
| Cinnamon bark | *Cinnamomum zeylanicum* | - | - | - | - | - | - |
| Oregano | *Origanum vulgare* | - | - | - | - | - | - |
| Thyme white | *Thymus zygis* | - | - | - | - | - | - |
| Lemongrass | *Cymbopogon flexuosus* | - | - | - | - | - | - |
| Bandit "Thieves" | Synergy blend | - | - | - | - | + | + |
| Sandalwood oil | *Santalum Spicatum* | - | - | - | - | + | + |
| Health shield | Synergy blend | - | - | + | - | + | + |
| Allspice | *Pimenta officinalis* | - | - | + | - | - | - |
| Amyris | *Amyris balsamifera* | - | - | + | - | - | - |
| Palmarosa | *Cymbopogon martinii* | - | - | + | - | - | - |
| Cinnamon leaf | *Cinnamomum zeylanicum* | - | - | + | + | + | + |
| Clove bud | *Eugenia caryophyllata* | - | - | + | + | + | + |
| Citronella | *Cymbopogon winterianus* | - | - | + | + | + | + |
| Geranium bourbon | *Pelargonium graveolens* | - | - | + | + | + | + |
| Marjoram (Sweet) | *Origanum marjorana* | - | - | + | + | + | + |
| Peppermint | *Mentha piperita* | - | - | + | + | + | + |
| Lemongrass | *Cymbopogon citratus* | - | - | + | + | + | + |
| Cornmint | *Mentha arvensis* | - | - | + | + | + | + |
| Elemi | *Canarium luzonicum* | - | - | + | + | + | + |
| Ho wood | *Cinnamomum camphora* | - | - | + | + | + | + |
| Head ease | Synergy blend | - | - | + | + | + | + |
| Lemon eucalyptus | *Eucalyptus citriodora* | - | - | + | + | + | + |
| Litsea cubeba | *Litsea cubeba* | - | - | + | + | + | + |
| Myrrh | *Commiphora myrrha* | - | - | + | + | + | + |
| Parsley seed | *Petroselinum sativum* | - | - | + | + | + | + |
| Coriander oil | *Coriandrum sativum* | - | - | + | + | + | + |
| Dillweed | *Anethum graveolens* | - | - | + | + | + | + |
| Hyssop | *Hyssopus officinalis* | - | - | + | + | + | + |
| Neroli | *Citrus aurantium* | - | - | + | + | + | + |
| Rosewood oil | *Aniba rosaeodora* | - | - | + | + | + | + |
| Tea Tree | *Melaleuca alternifolia* | + | - | + | + | + | + |
| Cajeput | *Melaleuca cajuputi* | + | - | + | + | + | + |
| Glove Bud | *Syzygium aromaticum* | + | - | + | + | + | + |
| Lavender | *Lavandula officinalis* | + | - | + | + | + | + |
| Sleep tight | Synergy blend | + | - | + | + | + | + |
| Vetiver | *Vetiveria zizanioides* | + | - | + | + | + | + |
| Palo Santo | *Bursera graveolens* | + | - | + | + | + | + |
| Sage oil | *Salvia officinalis* | + | - | + | + | + | + |
| Yarrow | *Achillea millefolium* | + | - | + | + | + | + |

<sup>a</sup> "EO" essential oil.
<sup>b</sup> “-” No obvious colonies grew on TSB plate after drug exposure; “+” Obvious colonies were found on TSB plate after drug exposure.

### MIC determination of the top active essential oils

We carried out antibiotic susceptibility testing to determine the activity of the top 10 active essential oils against growing *S. aureus*. As shown in Table 2, Oregano, Amyris and Sandalwood oil were the most active agents in inhibiting the growth of *S. aureus*, with the lowest MIC of 0.015% in our study. The growth of *S. aureus* was efficiently suppressed by Cinnamon bark at 0.03%. Allspice could inhibit the growth of *S. aureus* with an MIC of 0.06%, while Thyme white, Health shield, Bandit “Thieves”, Lemongrass (*Cymbopogon flexuosus*) and Palmarosa had the same MIC of 0.125% against *S. aureus*. Clinical drug gentamicin included as a control inhibited the growth of *S. aureus* with an MIC of 1 µg/mL.

**Table 2.**
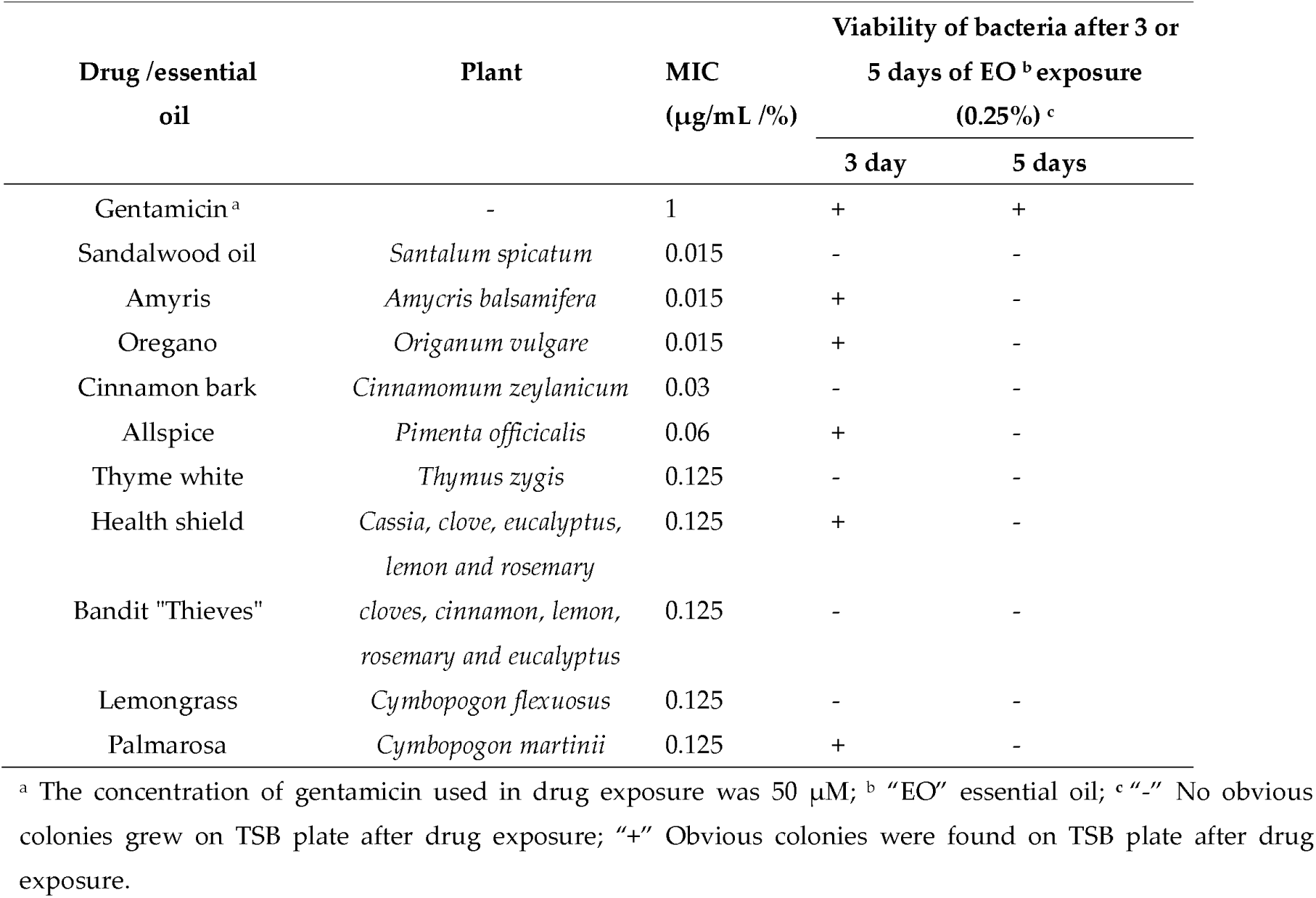
Activity of top 10 essential oils that are active against stationary phase *Staphylococcus aureus* in terms of their activity against growing bacteria (MIC) and non-growing bacteria in drug exposure.

| Drug /essential oil | Plant | MIC (µg/mL /%) | Viability of bacteria after 3 or 5 days of EO <sup>b</sup> exposure (0.25%) <sup>c</sup> |  |
| --- | --- | --- | --- | --- |
|  |  |  | 3 day | 5 days |
| Gentamicin <sup>a</sup> | - | 1 | + | + |
| Sandalwood oil | *Santalum spicatum* | 0.015 | - | - |
| Amyris | *Amycris balsamifera* | 0.015 | + | - |
| Oregano | *Origanum vulgare* | 0.015 | + | - |
| Cinnamon bark | *Cinnamomum zeylanicum* | 0.03 | - | - |
| Allspice | *Pimenta officinalis* | 0.06 | + | - |
| Thyme white | *Thymus zygis* | 0.125 | - | - |
| Health shield | *Cassia, clove, eucalyptus, lemon and rosemary* | 0.125 | + | - |
| Bandit "Thieves" | *cloves, cinnamon, lemon, rosemary and eucalyptus* | 0.125 | - | - |
| Lemongrass | *Cymbopogon flexuosus* | 0.125 | - | - |
| Palmarosa | *Cymbopogon martinii* | 0.125 | + | - |

<sup>a</sup> The concentration of gentamicin used in drug exposure was 50 µM; <sup>b</sup> "EO" essential oil; <sup>c</sup> "-" No obvious colonies grew on TSB plate after drug exposure; "+" Obvious colonies were found on TSB plate after drug exposure.

### Comparison of active essential oils in their ability to kill stationary phase *S. aureus*

We first tested the activity of tosufloxacin and other clinically used drugs against stationary phase *S. aureus* at 20 µM. As previously described [3], tosufloxacin could kill all stationary phase *S. aureus* cells after seven-day drug exposure, with no visible colonies remaining on TSB plate. Levofloxacin, ciprofloxacin and rifampin had weak activity with 10^4^∼10^5^ CFU/mL cells remaining after seven-day exposure. In contrast, other clinical drugs including linezolid, vancomycin, sulfamethoxazole, trimethoprim, azithromycin and gentamicin did not show obvious activity against stationary phase *S. aureus* even when the drug exposure was extended to seven days (Figure 1). In contrast, eight essential oils (Cinnamon bark, Oregano, Thyme white, Bandit “Thieves”, Lemongrass (*Cymbopogon flexuosus*), Health shield, Allspice, Palmarosa) at 0.25% concentration could eradicate all stationary phase cells after one-day exposure. Meanwhile, Amyris could clear all the cells after three-day exposure whereas Sandalwood oil could not wipe out the stationary phase *S. aureus* cells after seven-day exposure. At a lower concentration of 0.125%, we noticed that Oregano, Lemongrass (*Cymbopogon flexuosus*) and Thyme white still exhibited strong activity against stationary phase *S. aureus*, and no CFU could be detected after one-day exposure (Figure 2). Meanwhile, Cinnamon bark, Allspice, Amyris and Palmarosa could eradicate stationary phase *S. aureus* cells after three-day exposure. On the other hand, Bandit “Thieves”, Sandalwood oil and Health shield could not eradicate the stationary phase *S. aureus* culture even after seven-day exposure.

**Figure 1.**
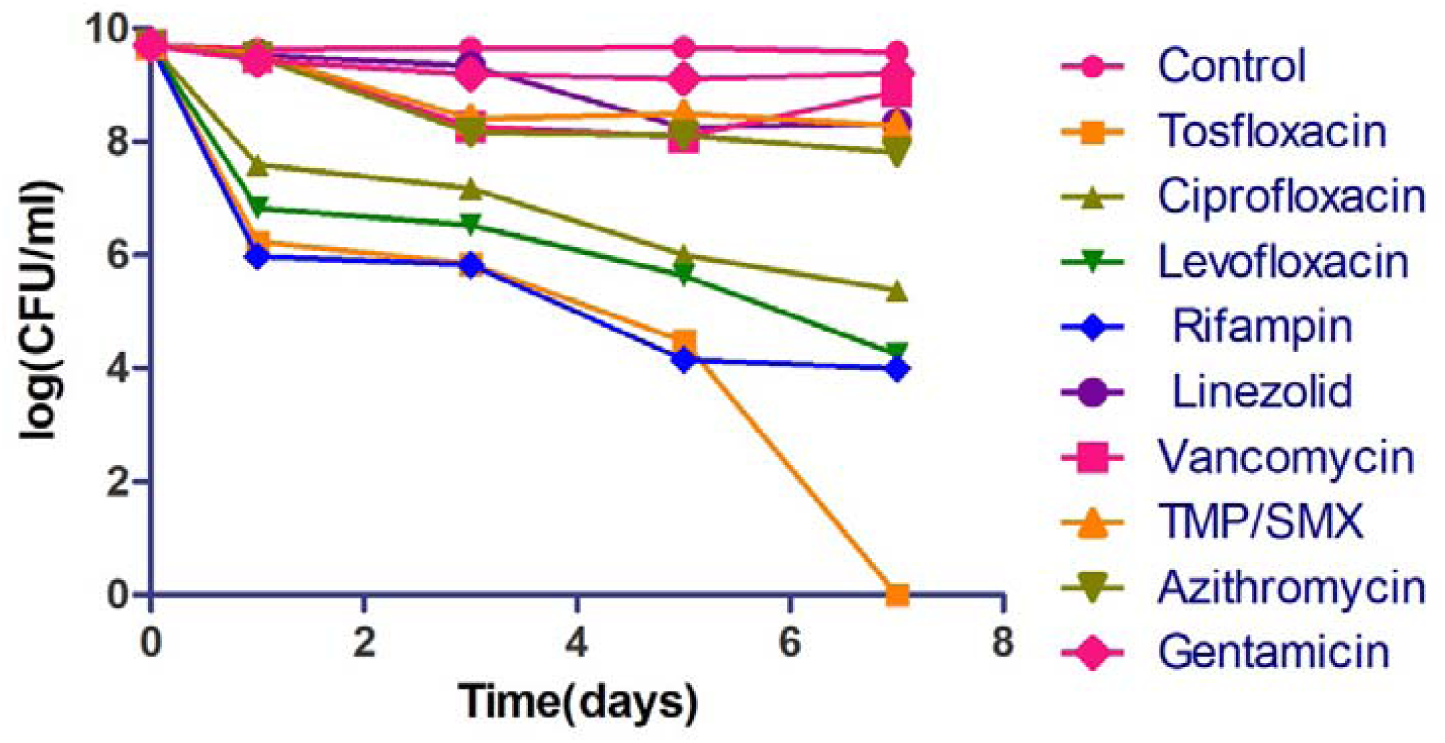
Activity of tosufloxacin and commonly used antibiotics against stationary phase *S. aureus*. Tosufloxacin had good anti-persister activity against *S. aureus*. Antibiotics commonly used to treat infections caused by *S. aureus* had poor activity against the stationary phase bacteria. The final concentration of antibiotics including tosufloxacin, ciprofloxacin, levofloxacin, rifampin, linezolid, vancomycin, sulfamethoxazole, trimethoprim, azithromycin and gentamicin, was all 20 µM. Sulfamethoxazole-trimethoprim is the combination of trimethoprim and sulfamethoxazole in a ratio of 5:1.

**Figure 2.**
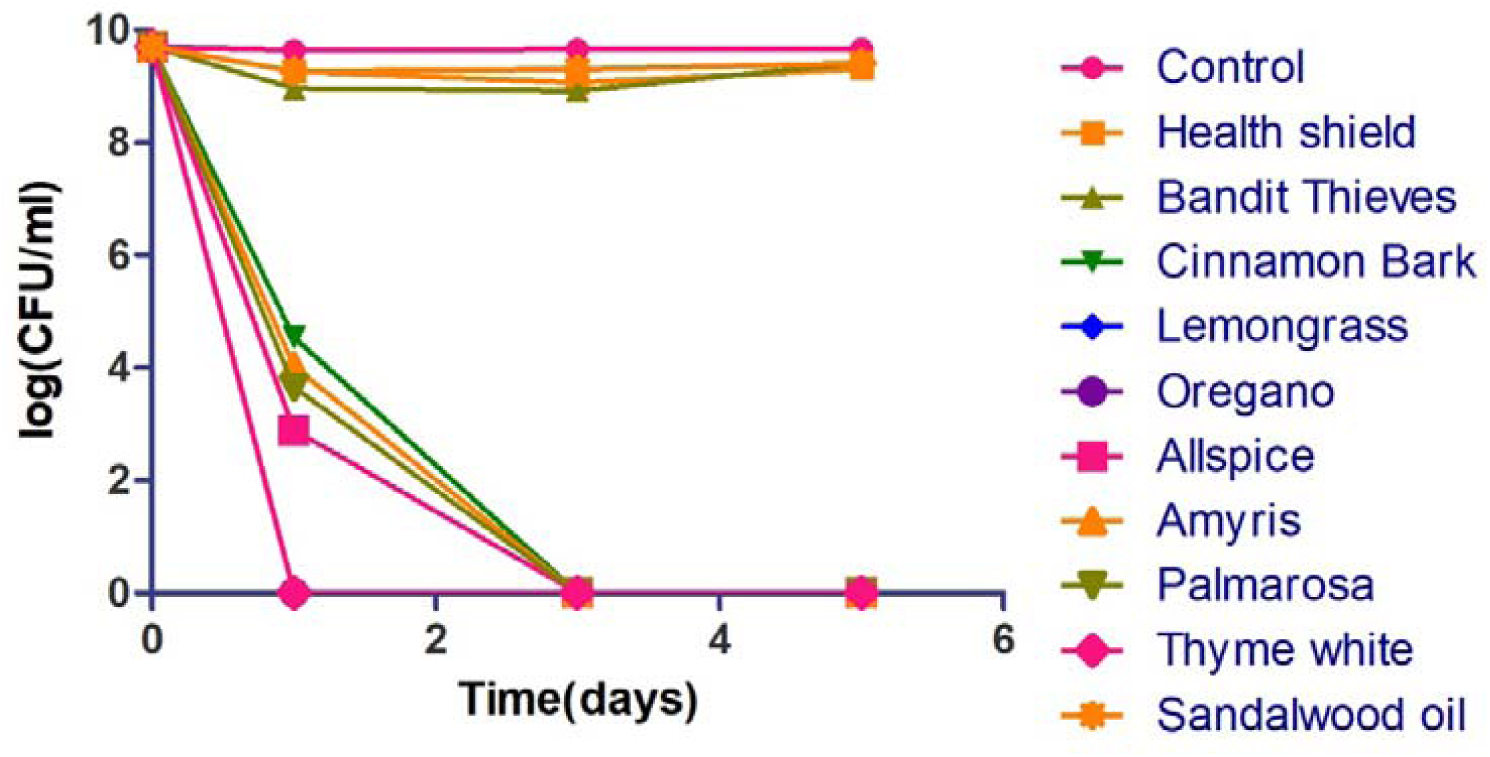
Activity of active essential oil candidates (0.125%) against stationary phase *S. aureus*. Oregano, Lemongrass (*Cymbopogon flexuosus*) and Thyme white could eradicate all stationary phase cells after one-day oil exposure. Cinnamon bark, Allspice, Amyris and Palmarosa could eradicate stationary phase *S. aureus* cells after three-day exposure. Bandit “Thieves”, Sandalwood oil and Health shield still could not wipe out the *S. aureus* stationary phase culture even after five-day exposure.

### Development of essential oil drug combinations to eradicate stationary phase *S. aureus* in vitro

It has been reported that synergistic activity between antibiotic and essential oil could occur, which achieved better bactericidal effect against growing *S. aureus* [12]. It is of great importance to include drugs that target persister bacteria in the treatment of infection diseases [4]. Based on our results, Oregano demonstrated high activity against not only log phase growing *S. aureus* with a low MIC but also stationary phase non-growing bacteria. Meanwhile, clinically used drugs had limited activity to kill *S. aureus* persisters. To more effectively eradicate the stationary phase *S. aureus*, we evaluated essential oil drug combinations using clinical drugs in combination with Oregano (0.025%). We found that some essential oil drug combinations were indeed much more effective than single drugs (Figure 3). Among them, rifampin + Oregano could completely eradicate all the stationary phase *S. aureus* after just one-day exposure. Tosufloxacin + Oregano could wipe out all stationary phase cells after three-day exposure. Meanwhile, levofloxacin + Oregano and ciprofloxacin + Oregano could kill all the stationary phase *S. aureus* after five-day exposure. These drug combinations showed much better activity than respective single drugs (10^4^ ∼ 10^6^ CFU/mL cells remaining) and somewhat better activity than single Oregano (10^4^ CFU/mL remaining). In contrast, other essential oil drug combinations such as linezolid + Oregano, vancomycin + Oregano, sulfamethoxazole + Oregano, trimethoprim + Oregano, azithromycin + Oregano and gentamicin + Oregano had limited activity against stationary phase cells, with 104 CFU/mL bacterial cells remaining even after five-day exposure, suggesting these combinations were not significantly better than Oregano alone (10^4^ CFU/mL remaining).

**Figure 3.**
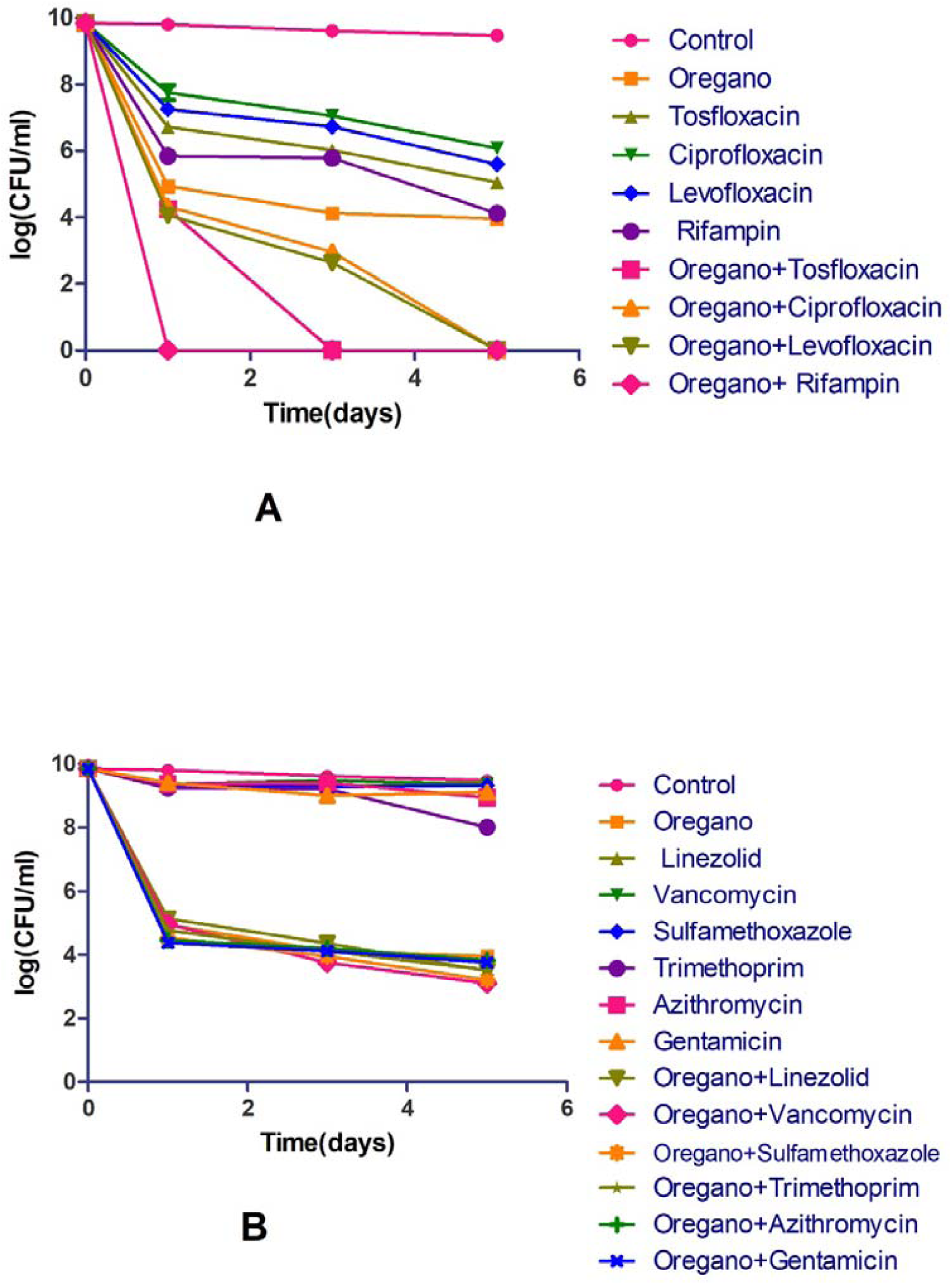
Comparison of the activity of Oregano in combination with different antibiotics against stationary phase *S. aureus*. Effects of ciprofloxacin, levofloxacin, tosufloxacin, rifampin alone and their combinations with Oregano are presented in (A). Effects of linezolid, vancomycin, sulfamethoxazole, trimethoprim, azithromycin, gentamicin and their combinations with Oregano are presented in (B). The final concentration of antibiotics is 5 µg/mL and the concentration of Oregano is 0.025%.

## Discussion

*S. aureus* is known to give rise to a diverse range of infections from mild skin infections to serious diseases such as endocarditis and osteomyelitis and biofilm infections. Persisters are dormant phenotypic variants of bacterial cells that are tolerant to antibiotics and genetically identical drug susceptible kin [2]. Since persisters were first identified in 1944 [13], there is considerable evidence that drug-tolerant persisters are the contributors to *S. aureus* persistent and relapsing infections [2, 14, 15], Meanwhile, treatment of persistent *S. aureus* infections has remained a challenge. It has been proposed to use persister drugs in the context of drug combination as in Yin-Yang model for more effective treatment of persistent infections [16]. Although previous study has screened FDA-approved drug library to identify agents that have good activity against stationary phase *S. aureus* [3], only few useful hits such as tosufloxacin and clinafloaxin were identified. Although ADEP4, an experimental acyldepsipeptide antibiotic killing *S. aureus* persisters in combination with rifampin has been reported to cure a deep wound infection in a mouse model [17], its validity in treating other persistent infections in other disease models remains to be confirmed. Thus, while it is of great importance to include drugs targeting persister bacteria in the treatment of *S. aureus* infections, the choice of persister drugs that may be useful is quite few. Since most studies of essential oil activity on *S. aureus* were performed on log phase growing bacteria [6, 7], here we set out to determine the activity of a large panel of essential oils against stationary phase *S. aureus* cultures enriched in persister bacteria. Interestingly, we identified a range of essential oils that have strong activity against stationary phase cultures of *S. aureus* that may be useful for more effective treatment of persistent *S. aureus* infections.

Essential oils, a widely studied alternative against antibiotic resistant bacteria, are concentrated volatile liquids extracted from plants. While there are some reports on activity of essential oils against log phase *S. aureus*, the number of evaluated essential oils is small (just one or two kinds of essential oils) [6, 8], and their activity against stationary phase *S. aureus* cultures has not been studied [6, 8]. In this study, we evaluated a panel of 143 essential oils for their activity against stationary phase *S. aureus.* We identified 9 essential oils (at 0.25% concentration) that are more active than persister drug tosufloxacin (20 µM), a quinolone drug control that could eradicate stationary phase *S. aureus*. Among them, 7 essential oils (Cinnamon bark, Oregano, Thyme white, Lemongrass (*Cymbopogon flexuosus*), Allspice, Amyris, Palmarosa) showed outstanding activity against stationary phase *S. aureus* at 0.125% concentration (Figure 2). Meanwhile, all of the top 9 essential oils showed high activity against growing *S. aureus* (Table 2), of which some essential oils such as Thyme white, Oregano and Cinnamon bark have been reported to have high activity against log phase *S. aureus* in previous studies [7, 18, 19], but Lemongrass (*Cymbopogon flexuosus*), Bandit “Thieves”, Health shield, Allspice, Amyris, Palmarosa were first reported in this study. Compared with our previous work on activity of essential oils against stationary phase *E. coli*, some essential oils including Cinnamon bark, Oregano, Bandit “Thieves”, Health shield and Allspice exhibited outstanding activity against both Gram-positive *S. aureus* and Gram-negative *E. coli* [20], while it seems that Thyme white, Lemongrass (*Cymbopogon flexuosus*), Amyris, Palmarosa just showed high activity against *S. aureus* [20], while Cinnamon leaf, Clove bud and *Syzygium aromaticum* were only active against *E. coli* [20]. Moreover, although some studies indicate that certain active essential oils including their main active components such as carvacrol or eugenol could induce membrane damage by causing loss of cellular contents [9, 21, 22], there are limited studies available that focus on the active components and the mechanisms of antimicrobial action of essential oils in general. Here, we identified some active essential oils against stationary phase *S. aureus.* Further studies are needed to determine the main active components and the mechanisms of action of the active essential oils identified in this study.

Oregano is known to be one of the most effective essential oils against a wide variety of pathogens, including *Pseudomonas sp*., *Salmonella sp*., *Escherichia coli* and *Borrelia burgdorferi* [7, 23]. In this study, Oregano exhibited high activity against not only log phase growing *S. aureus* with a low MIC of 0.015% but also stationary phase non-growing bacteria with complete clearance without any regrowth at 0.125% concentration. Remarkably, when combined with some currently recommended antibiotics for *S. aureus* infections, Oregano showed a positive enhancement effect in increasing the activity of some antibiotics (quinolones, rifampin) against stationary phase *S. aureus* (Figure 3). When combined with rifampin, the combination showed outstanding activity with 100% clearance after just one-day exposure. When combined with tosufloxacin and two other quinolone drugs (levofloxacin and ciprofloxacin), the combinations could wipe out all stationary phase cells after three-day or five-day exposure. The synergistic effect of Oregano and the four drugs may have implications for improved treatment of *S. aureus* persistent infections. Further studies should be carried out to confirm if such combination approaches are useful in animal models.

Additionally, we found Cinnamon bark, Thyme white, Lemongrass (*Cymbopogon flexuosus*), Allspice, Amyris and Palmarosa showed excellent activity against stationary phase *S. aureus* at a low concentration of 0.125% (Figure 2). Cinnamon bark was reported to have activity against bacteria, fungi, inflammation, cancer and diabetes [24]. Also, it was demonstrated to be a potential antibacterial agent that exhibited strong activity against methicillin resistant *S. aureus* (MRSA) [25]. In this study, Cinnamon bark also showed its remarkable activity against non-growing stationary phase *S. aureus*. Thyme white, extracted from *Thymus zygis*, showed great activity against both log phase and stationary phase *S. aureus* at the same concentration of 0.125%. Essential oils obtained from *Thymus* species were often compared for their antibacterial and antioxidant activity. And oil from *Thymus zygis* was the most active one towards log phase Gram-positive and Gram-negative bacteria [18, 26]. Our results highlighted its antibacterial activity not only against growing *S. aureus* bacteria but also non-growing stationary phase cells. Lemongrass from two different plants (*Cymbopogon flexuosus* and *Cymbopogon citratus*) were evaluated in this study. While Lemongrass from *Cymbopogon flexuosus* could kill all the stationary phase *S. aureus* in just one day at 0.125%, Lemongrass from *Cymbopogon citratus* showed obvious activity only at a high concentration (0.5%). This provides the basis for further testing of *Cymbopogon flexuosus* in animal models of infection. Allspice is widely known as a popular spice in food processing [27], here, its activity against *S. aureus* may facilitate its usage for antibacterial purpose. Compared with other essential oils, there are few studies discussing bioactivity of Amyris. One study revealed that vapor delivery of Amyris could alter pyrethroid efficacy and detoxification enzyme activity in mosquitoes [28]. Our new finding of Amyris activity on *S. aureus* in this study may contribute to more bioactivity of Amyris and therapeutic use of *Amycris balsamifera.* Palmarosa, known as *Cymbopogon martini*, is used in Ayurvedic medicine to relieve nerve pain for skin problems and as a skin tonic in aromatherapy due to its antimicrobial properties [29]. While its immunomodulatory activity is based on geraniol [29], the main component of Palmarosa active against stationary phase *S. aureus* is unknown and will be determined in the future.

Along with the essential oils which showed strong activity against *S. aureus* persisters at 0.125%, there were three essential oils (Bandit “Thieves”, Sandalwood oil and Health shield) that showed obvious activity only at higher concentrations. Bandit “Thieves” and Health shield could eradicate all stationary phase cells after one-day exposure at 0.25% concentration. Sandalwood oil exhibited obvious activity at 0.5% concentration. Both Bandit “Thieves” and Health shield are synergy blend of essential oils. While Bandit “Thieves” contains clove, cinnamon, lemon, rosemary and eucalyptus oils, Health shield is a mixture of essential oils from cassia, clove, eucalyptus, lemon and rosemary. They were active against the growing of *S. aureus* with the same MIC value of 0.125% and showed similar activity against stationary phase *S. aureus* cells (Table 2 and Figure 2). Sandalwood oil in this study is obtained from *Santalum spicatum* (Australian Sandalwood). There are two other kinds of Sandalwood oil: East Indian Sandalwood oil extracted from *Santalum album* and New Caledonian Sandalwood oil prepared from the wood of *Santalum austrocaledonicum* [30]. Sandalwood oil from East Indian is widely studied as an attractive natural therapeutic for inflammatory skin diseases [31]. On the other hand, Sandalwood oil from *Santalum spicatum* has high commercial value for applications in aromatherapy and for the production of cosmetics such as soaps, creams and powder [30]. In this study, Sandalwood oil extracted from *Santalum spicatum* showed high activity against *S. aureus*, which demonstrated that it may be a promising antibacterial agent.

In summary, this is the first study of a large collection of 143 essential oils for activity against stationary phase *S. aureus* where we identified several promising essential oils. The top hits are Oregano, Cinnamon bark, Thyme white, Lemongrass (*Cymbopogon flexuosus*), Bandit “Thieves”, Sandalwood oil, Health shield, Allspice, Amyris, Palmarosa. Meanwhile, we found in drug combination study with essential oil (Oregano) and antibiotics that some potent combinations such as Oregano plus quinolones or rifampin could effectively eradicate *S. aureus* persisters in vitro. Further studies should be carried out to identify the active components, evaluate safety, pharmacokinetics, and their activity to eradicate *S. aureus* persistent infections in animal models.

## Funding

We acknowledge the support by the Einstein-Sim Family Charitable Fund and Guangci Distinguished Young Scholars Program (GCQN-2017-C11).

